# A mechanism for prime-realignment during influenza A virus replication

**DOI:** 10.1101/138487

**Authors:** Judith Oymans, Aartjan J.W. te Velthuis

## Abstract

The influenza A virus genome consists of eight segments of single-stranded RNA. These segments are replicated and transcribed by a viral RNA-dependent RNA polymerase (RdRp) that is made up of the influenza virus proteins PB1, PB2 and PA. To copy the viral RNA (vRNA) genome segments and the complementary RNA (cRNA) segments, the replicative intermediate of viral replication, the RdRp must use two promoters and two different *de novo* initiation mechanisms. On the vRNA promoter, the RdRp initiates on the 3’ terminus, while on the cRNA promoter the RdRp initiates internally and subsequently realigns the nascent vRNA product to ensure that the template is copied in full. In particular the latter process, which is also used by other RNA viruses, is not understood. Here we provide mechanistic insight into prime-realignment during influenza virus replication and show that it is controlled by the priming loop and a helix-loop-helix motif of the PB1 subunit of the RdRp. Overall, these observations advance our understanding of how the influenza A virus initiates viral replication and amplifies the genome correctly.

**Importance:** Influenza A viruses cause severe disease in humans and are considered a major threat to our economy and health. The viruses replicate and transcribe their genome using an enzyme called the RNA polymerases. To ensure that the genome is amplified faithfully and abundant viral mRNAs are made for viral protein synthesis, the RNA polymerase must work correctly. In this report, we provide insight into the mechanism that the RNA polymerase employs to ensure that the viral genome is copied correctly.

## Introduction

The influenza A virus (IAV) is an important pathogen that causes seasonal epidemics and occasional pandemics in humans. The IAV genome comprises eight segments of single-stranded, negative sense viral RNA (vRNA) that exist in the context of viral ribonucleoproteins (vRNPs (1). These vRNPs consist of one vRNA segment, a copy of the viral RNA-dependent RNA polymerase (RdRp), and a helical coil of the viral nucleoprotein (NP) and during IAV infections the vRNPs are released from viral particles and imported into the nucleus of the host cell. In the nucleus, the vRNPs replicate the vRNAs via a complementary RNA (cRNA) intermediate and they transcribe the vRNAs to form viral mRNAs (1, 2). The latter molecules are exported from the nucleus and translated by cellular ribosomes, while cRNAs are bound by new NP and RdRp molecules in order to form cRNPs capable of synthesising new vRNAs (3).

Both IAV replication and transcription are catalysed by the RdRp (1, 2). This RdRp is a 250 kDa heterotrimer that consists of the viral proteins polymerase basic protein 1 (PB1), polymerase basic protein 2 (PB2), and polymerase acidic protein (PA) (1, 4). The PB1 subunit, the N-terminal third of PB2 and the C-terminal two-thirds of PA form the conserved RdRp domain (5–7), while the remaining parts of PA and PB2 form flexible domains at the periphery of the polymerase core (Fig 1A). These domains are important for cleaving cellular host mRNAs, a process that yields capped RNA primers that are essential for viral transcription (1, 4).

**Figure 1.**
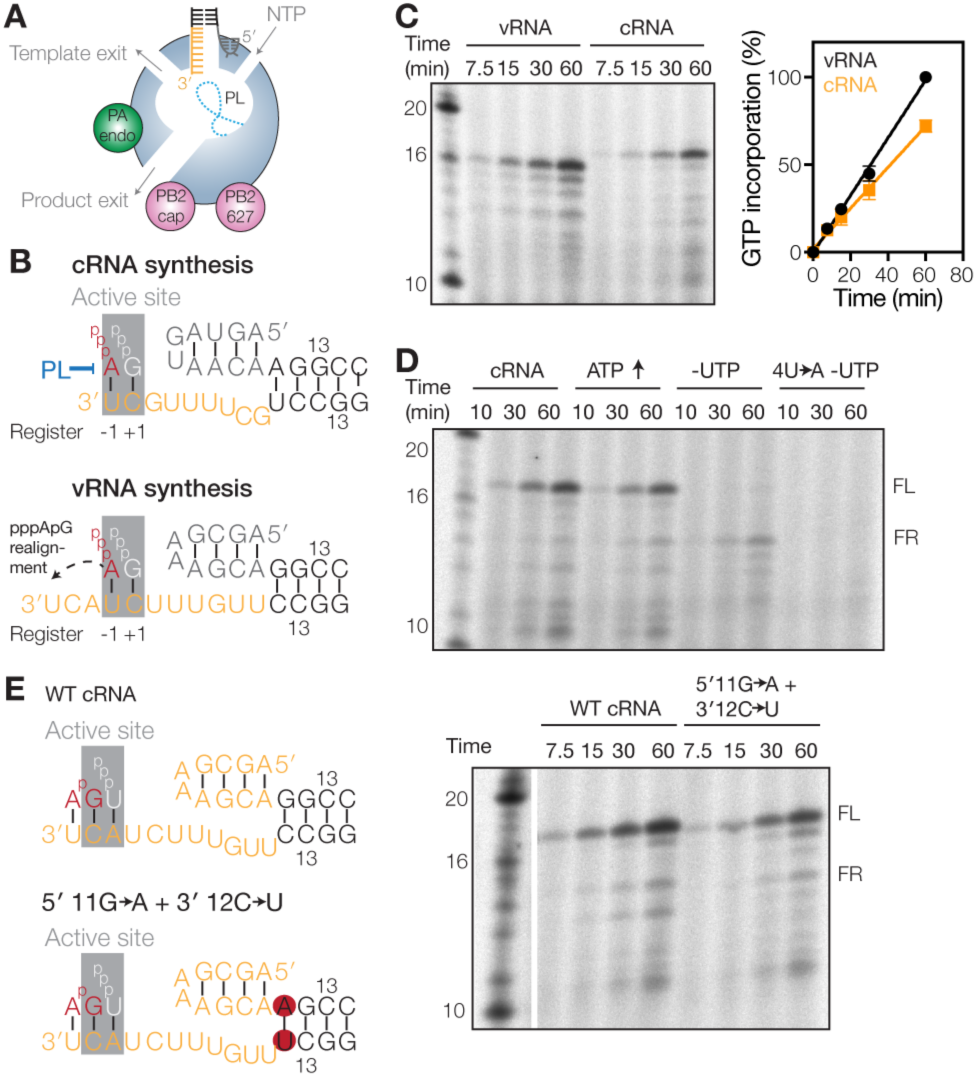
Failed prime-realign events can be detected *in vitro*. (**A**) Model of the influenza A virus RdRp. The core of the RdRp is shaded blue, the PA endonuclease (PA endo) green, and the PB2 cap binding domain (PB2 cap) and 627 domain (PB2 627) pink. The priming loop (PL) is indicated as a dotted line. (**B**) Schematics of IAV initiation during cRNA and vRNA synthesis. The priming loop (PL) is indicated in blue. The active site (grey) positions −1 and +1 are indicated below each schematic. (**C**) Time course of ApG extension on a vRNA or cRNA promoter. The graph shows the percentage of [α-^32^P]GTP incorporation relative to the activity on the vRNA promoter. Error bars indicate standard deviation (*n* = 3). (**D**) Time course of ApG extension on a wild-type or 4U→A mutant cRNA promoter. The addition of 0.5 mM ATP or the omission of UTP is indicated. (**E**) Schematic of ApG extension on a wild-type cRNA promoter or a promoter where the first G-C base-pair of the promoter duplex was mutated to A-U (5ʹ 11G→A + 3ʹ 12C→U). Gel image shows ApG extension assay on the wild-type cRNA promoter or the 5ʹ 11G→A + 3ʹ 12C→U promoter mutant.

The viral RdRp binds the 5ʹ and 3ʹ terminal ends of the vRNA or cRNA, also called the vRNA or cRNA promoters respectively (8–10), via a binding pocket above the NTP channel (5, 6, 11). The 3ʹ end of either promoter can translocate to the active site (12) (Fig. 1A), allowing *de novo* initiation on the terminal 3ʹ UC of the vRNA (Fig. 1B) or *de novo* initiation at positions 4U and 5C of the 3ʹ terminus of the cRNA (Fig. 1B) (13, 14). However, the two *de novo* initiation mechanisms are markedly different. Terminal *de novo* initiation on a vRNA promoter, but not internal *de novo* initiation on a cRNA promoter, critically depends on the PB1 priming loop (Fig. 1B) (13). By contrast, internal initiation requires realignment of the pppApG initiation product to bases 3ʹ 1U and 2C of the cRNA prior to elongation (13, 14) (Fig. 1B). A failure to perform this realignment step would generate a vRNA lacking the first 3 nucleotides of the canonical vRNA 5ʹ terminus. Since the vRNA 5ʹ terminus is part of the vRNA promoter and critical for the activity and conformational changes of the influenza virus RdRp (5–7), such a truncated vRNA would likely not support efficient cRNA and mRNA synthesis. Current evidence suggests that a number of negative strand RNA viruses use a similar or related realignment process to ensure faithful replication of their genome (14–16), but the molecular mechanism that controls these realignment processes is currently not understood.

In this study, we investigated the prime-realign mechanism using a combination of cell-based RNP reconstitution assays, structure-guided mutagenesis, and *in vitro* activity assays. We show that the IAV RdRp uses its priming loop to enforce prime-realignment and thus correct vRNA synthesis during replication. Our observations provide novel mechanistic insight into IAV RNA synthesis and expand our current view of the role of the priming loop.

## Results

### Prime-realignment is essential for viral RNA synthesis

The IAV RdRp catalyses mRNA and cRNA synthesis from the vRNA promoter and vRNA synthesis from the cRNA promoter. We previously showed that the initiation activity of the IAV RdRp on these two promoters is similar during viral replication (13). However, a side-by-side comparison of the RdRp’s extension activity on these two promoters shows that the RdRp activity is ~30% weaker on the cRNA promoter than on the vRNA promoter (Fig. 1C; note that the full-length products of the vRNA and cRNA promoter, a 14 nt and 15 nt product respectively, migrate near the 16-nt and 17-nt marker bands, because the ApG that primes the reaction lacks a 5ʹ phosphate, which reduces the negative charge and migration of the viral products relative to the 5ʹ phosphorylated marker (13)). A similar observation was recently made for the activities of the influenza B virus RdRp on the influenza B virus vRNA and cRNA promoters (17). This difference in promoter activity may be explained, at least in part, by the prime-realign mechanism that the RdRp uses during initiation on the cRNA promoter (Fig. 1B) or the difference in affinity of the RNA polymerase for the two templates (12). So far, the efficiency of the former and the mechanism behind the realignment process has not been studied in detail.

To study how the IAV RdRp coordinates prime-realignment, we attempted to measure the formation of failed realignment (FR) products in ApG extension assays using influenza A/WSN/33 virus RdRp preparations. We decided to use the fact that ApG extension after a realignment step requires UTP incorporation, while a failed realignment event utilises ATP for extension (Fig. 1B). To favour the latter event, we either doubled the ATP concentration or omitted UTP from the reaction. Only under the UTP-free condition, we observed a loss of the 15 nt full-length (FL) product and the appearance of a 12 nt main product (Fig. 1D). Moreover, this 12 nt product was not synthesised in reactions where 4U of the 3ʹ cRNA strand was mutated to A (4U→A) to prevent ApG priming at position 4/5 and reduce internal elongation (Fig. 1D), confirming that the FR product was dependent on internal ApG extension.

To investigate whether the amount of FR product was dependent on the stability of the model promoter duplex, we replaced the first G-C base pair of the duplex of the cRNA promoter with an A-U base pair to make the duplex less stable (Fig. 1E). No increase in FR product formation was observed in reactions containing the mutant cRNA promoter duplex (Fig. 1E), suggesting that the duplex of the model cRNA promoter does not influence the prime-realignment mechanism. Together these observations thus suggest that failed prime-realignment events can be detected and identified *in vitro*, but that they do not occur frequently. This implies that the efficiency of prime-realignment is relatively high and that the difference in promoter activity (Fig. 1C) may be largely explained by the different binding efficiencies of the RdRp to the two viral promoters (12).

### PB1 V273 modulates prime-realignment during vRNA synthesis

Realignment of the pppApG initiation product during vRNA synthesis is likely controlled by the RdRp structure. Inside the RdRp, the nascent A-form duplex, consisting of template and product RNA, is guided away from the active site by a conserved helix-turn-helix structure (Fig. 2A). To investigate whether residues in this structure contributed to the prime-realignment mechanism, we engineered alanine substitutions of conserved PB1 residues S269, L271, P272 or V273 at the top of the helix-turn-helix (Fig. 2B), creating S269A, L271A, P272A and V273A, respectively. A PB1 mutant containing alanine substitutions of two critical active site aspartates (PB1 DD445-446AA; PB1a) (18) was used as negative control. Using IgG sepharose purification followed by SDS-PAGE analysis and silver-staining or Western blot we verified that the mutations had no effect on heterotrimer formation (Fig. 2C).

**Figure 2.**
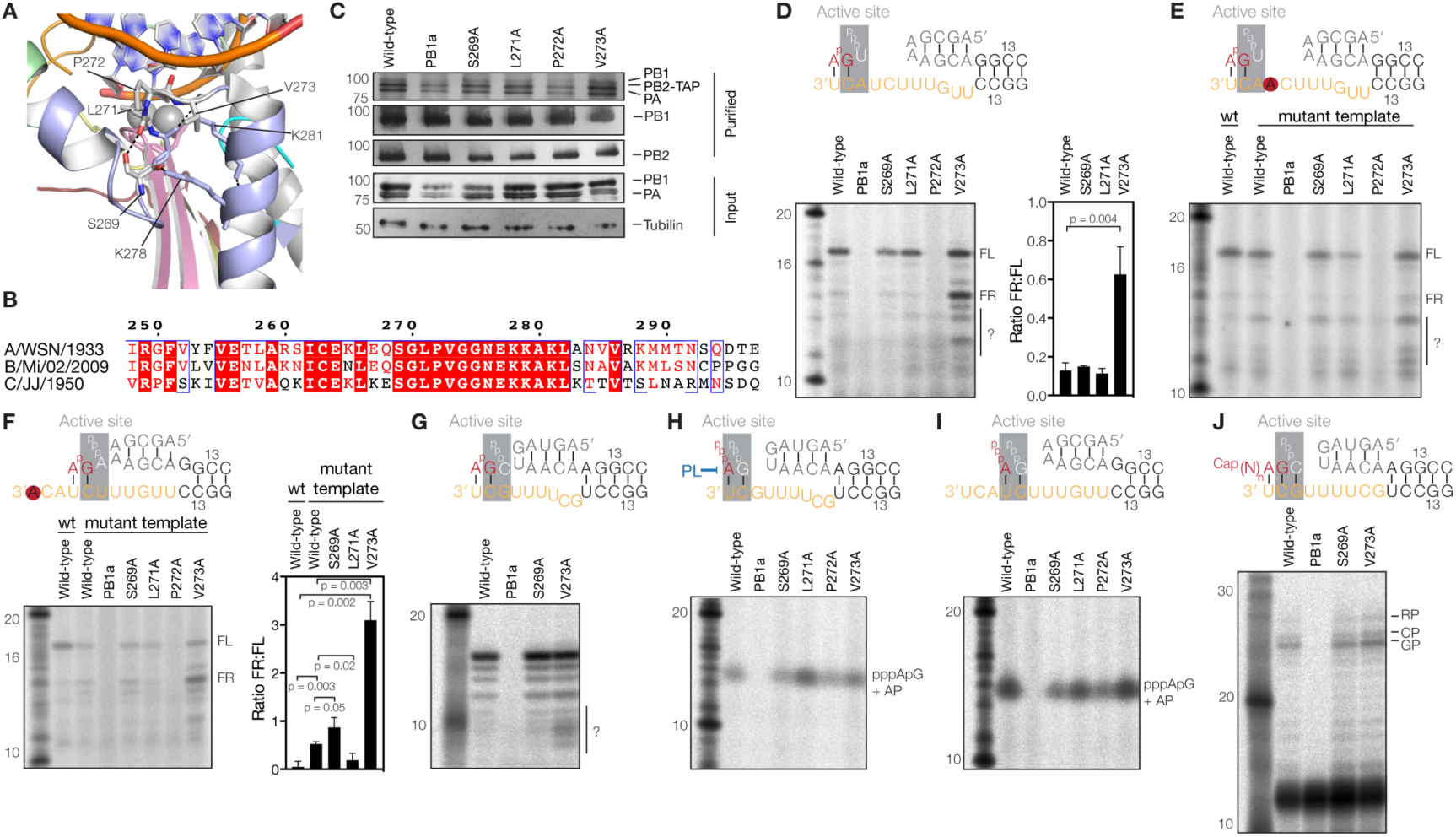
PB1 V273 affects prime-realignment. (**A**) Superposed structure of the bat influenza A virus RdRp (PDB 4WSB) with the poliovirus 3D^pol^ RdRp elongation complex (PDB 3OL7). For the 3D^pol^ complex, the template strand, nascent strand, and magnesium ions are shown. For the bat influenza A virus RdRp, the PB1 subunit is shown in light blue, with polymerase motifs A, C, D, and F coloured yellow, pink, red, and pale green, respectively. Polar interactions between amino acids of the helix-loop-helix structure are indicated with dotted lines. Additional side-chains are shown for reference. (**B**) Amino acid alignment of the PB1 helix-turn-helix structure of the palm subdomain. PB1 sequences of the influenza A virus A/WSN/33 (H1N1), influenza B virus B/Michigan/22687/09, and influenza C virus C/JJ/50 are shown. Identical residues are shaded red and conserved residues are surrounded with blue boxes. Secondary structure annotations are based on PDB 4WSB. (**C**) SDS-PAGE and silver staining of recombinant influenza A virus RdRps isolated from 293T cells or Western-blot analysis of PB1 and PB2 in isolated RdRps. Bottom panel shows expression of PB1 and PA RdRp subunits in 293T cells. Also a tubulin loading control is shown. (**D**) ApG extension on a cRNA promoter. The graph shows the ratio between the quantified FR and FL signals of three independently purified RdRp sets. The p-value was determined using an unpaired t-test. Unknown products are indicated with a question mark. (**E**) ApG extension on the 4U→A mutant cRNA promoter. Question mark indicates an increase in unknown RNA products. (**F**) ApG extension on the 1U→A mutant cRNA promoter. The graph shows the mean FR to FL product ratio of three independently purified RdRp sets. The p-values were determined using an unpaired t-test. (**G**) ApG extension on a vRNA promoter. Unknown products are indicated with a question mark. In each graph, the error bars indicate standard deviation (*n* = 3). (**H**) Terminal pppApG synthesis on a cRNA promoter in the presence of ATP and [α-^32^P]GTP. The reactions were treated with alkaline phosphatase (AP) to better separate the radioactive product from the non-incorporated [α-^32^P]GTP and free phosphates. (**I**) Internal pppApG synthesis on a vRNA promoter. (**J**) Extension of a radiolabelled capped 11-nucleotide long RNA primer ending in 3ʹ AG. This extension reaction yields a product initiated at G3 (GP) or a product initiated at 2C (CP) of the vRNA promoter and an additional realignment product (RP) (39).

To investigate the effect of the mutations on the realignment efficiency, we performed ApG extension assays on a cRNA promoter and analysed the reactions by 20% denaturing PAGE. No change in FR product formation was observed in reactions containing PB1 mutants S269A and L271A (Fig. 2D). By contrast, mutant P272A failed to extend the ApG dinucleotide (Fig. 2D), while mutant V273A produced significantly more FR than wild-type (Fig. 2D). In addition, V273A produced RNAs migrating faster than the FR product, but the nature of these RNA species is presently unknown. To confirm that the V273A FR product had been produced through internal elongation, we measured the activity of V273A on the 4U→A mutant promoter to reduce internal ApG binding and observed a substantial reduction in the FR signal without impairment of FL production (Fig. 2E). In support of this observation, mutation of 1U of the cRNA 3ʹ strand to A (1U→A), which reduces ApG priming at position 1U/2C, reduced FL product formation for all RdRps tested and increased the FR:FL ratio ~3-fold for V273A (Fig. 2F). To verify that the V273A mutation was specifically affecting the prime-realignment mechanisms and that no other RdRp activities were affected, we performed ApG extensions on the vRNA promoter (Fig. 2G), and *de novo* initiation and transcription initiation assays (Fig. 2H-J). We found no effect of the V273A mutation on the polymerase activity in any of these assays. By contrast, mutations S269A, L271A and P272A had small effects on *de novo* initiation (Fig. 2H-J). Overall, these observations suggest that V273 at the tip of the PB1 helix-turn-helix primarily affects the prime-realign process.

We next investigated the activity of V273 in cell culture. To this end, we used a mini replicon assay that relies on the reconstitution of vRNPs from plasmid-expressed IAV RdRp subunits, NP and a segment 6 (neuraminidase (NA)-encoding) vRNA template (Fig. 3). After RNA extraction, the synthesis of the viral RNA species (vRNA, cRNA and mRNA) was measured using primer extensions of 5ʹ radiolabelled primers (Table 1). As shown in Fig. 3, viral RNA synthesis by V273A, but also S269A, was impaired compared to wild-type and found to have a differential effect on cRNA and mRNA synthesis. These observations corroborate previous RNP reconstitutions with the same mutants (19) and experiments showing that mutations V273A, V273L or V273D reduce the fitness of A/WSN/33 viruses by >90% (20). Based on these observations, we suggest that inefficient realignment of the nascent vRNA results in the synthesis of truncated vRNA products that may not be bound by the IAV RdRp. In turn, these truncated vRNA may be degraded by the host cell, which restricts viral cRNA and mRNA synthesis to primary RNA synthesis and reduces viral RNA levels without directly affecting the mechanism of transcription or cRNA synthesis. Overall, these results imply that correct realignment and/or internal initiation is essential for influenza virus replication and transcription and that this process is modulated, at least in part, by PB1 residue V273.

**Figure 3.**
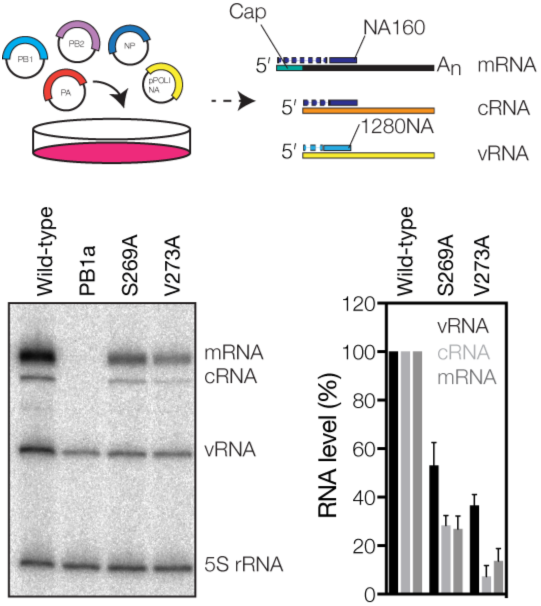
PB1 V273 affects RNA synthesis in cell culture. Schematic of the RNP reconstitution assay and analysis of the steady state segment 6 RNA levels. The graph shows the mean RNA levels of three independent experiments after subtraction of the PB1 active site control signal. Error bars indicate standard deviation (*n* = 3).

**Table 1:**
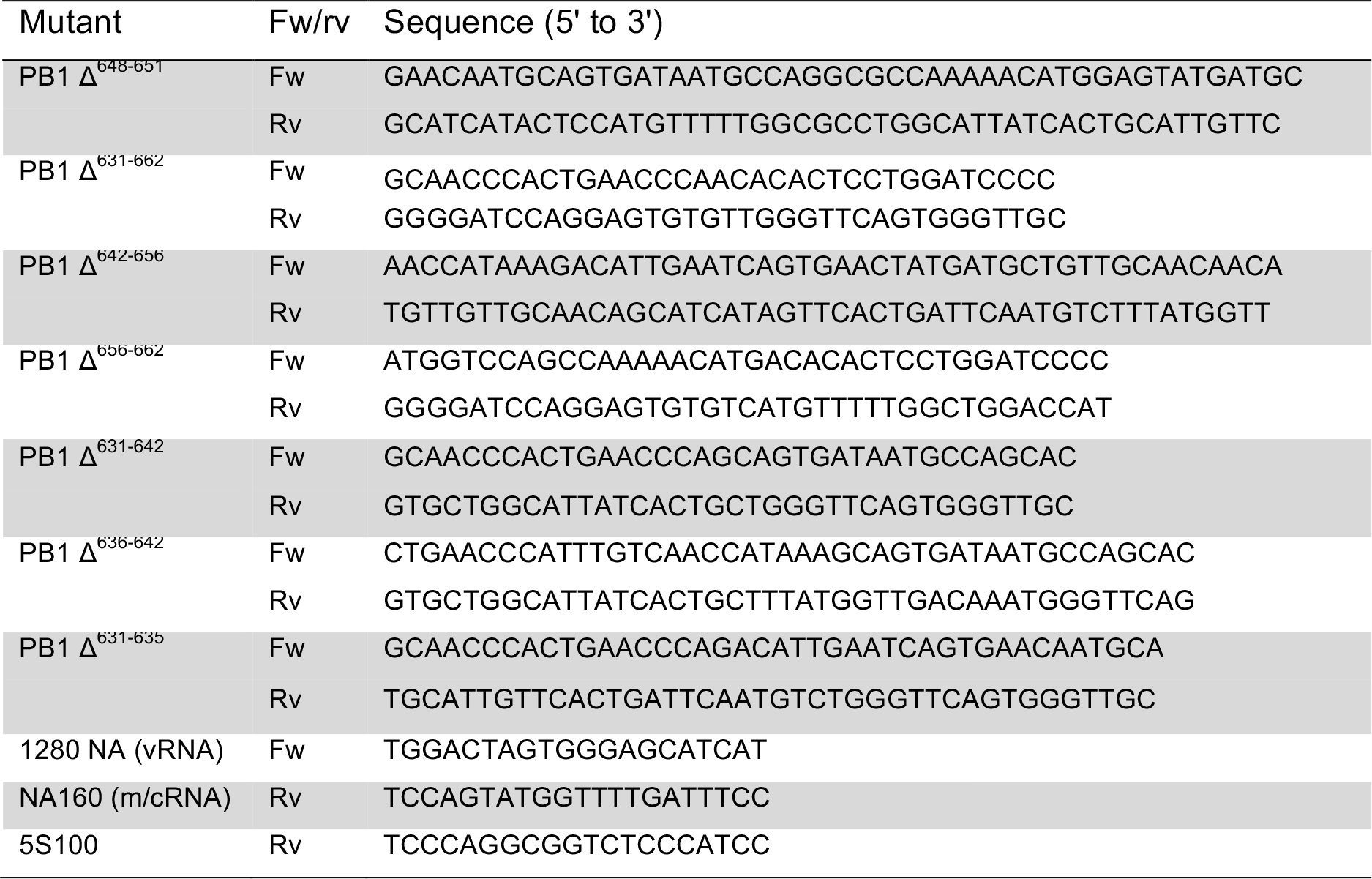
Primers used for PB1 mutagenesis and primer extension.

### The priming loop is important for viral RNA synthesis in cell culture

Research of other RNA virus RdRps has shown that the correct positioning of the template in the active site is dependent on the priming loop (21–23). In the IAV RdRp, the priming loop resides downstream of the active site (Fig. 4A), above the helix-turn-helix containing PB1 V273. To investigate whether the priming loop played a role in prime-realignment, we engineered seven deletions in the influenza A/WSN/33 (H1N1) virus PB1 subunit based on the different conformations of the priming loop in current crystal structures (Fig. 4B) and the sequence conservation of the priming loop among influenza A, B and C viruses (Fig. 4C). The deleted sections (a-e) are indicated on the crystal structure of the bat influenza A virus RdRp and a PB1 sequence alignment (Fig. 4B and 4C). In RNP reconstitutions, all mutants were significantly impaired in viral RNA synthesis (Fig. 4D). Due to the interdependence of viral replication and transcription in cell culture (i.e., some amplification of the template vRNA is required to observe mRNA signals above background), this result was expected for mutants Δ648-651, Δ642-656 and Δ631-662, because they all lack the tip of the priming loop that is critical for the initiation of cRNA synthesis (13). Indeed, none of these mutations was compatible with virus growth. The impaired activity of mutants Δ656-662, Δ631-642, Δ636-642 and Δ631-635 suggests that also the middle as well as the N-and C-terminal anchor points of the priming loop play an important role in RdRp activity.

**Figure 4.**
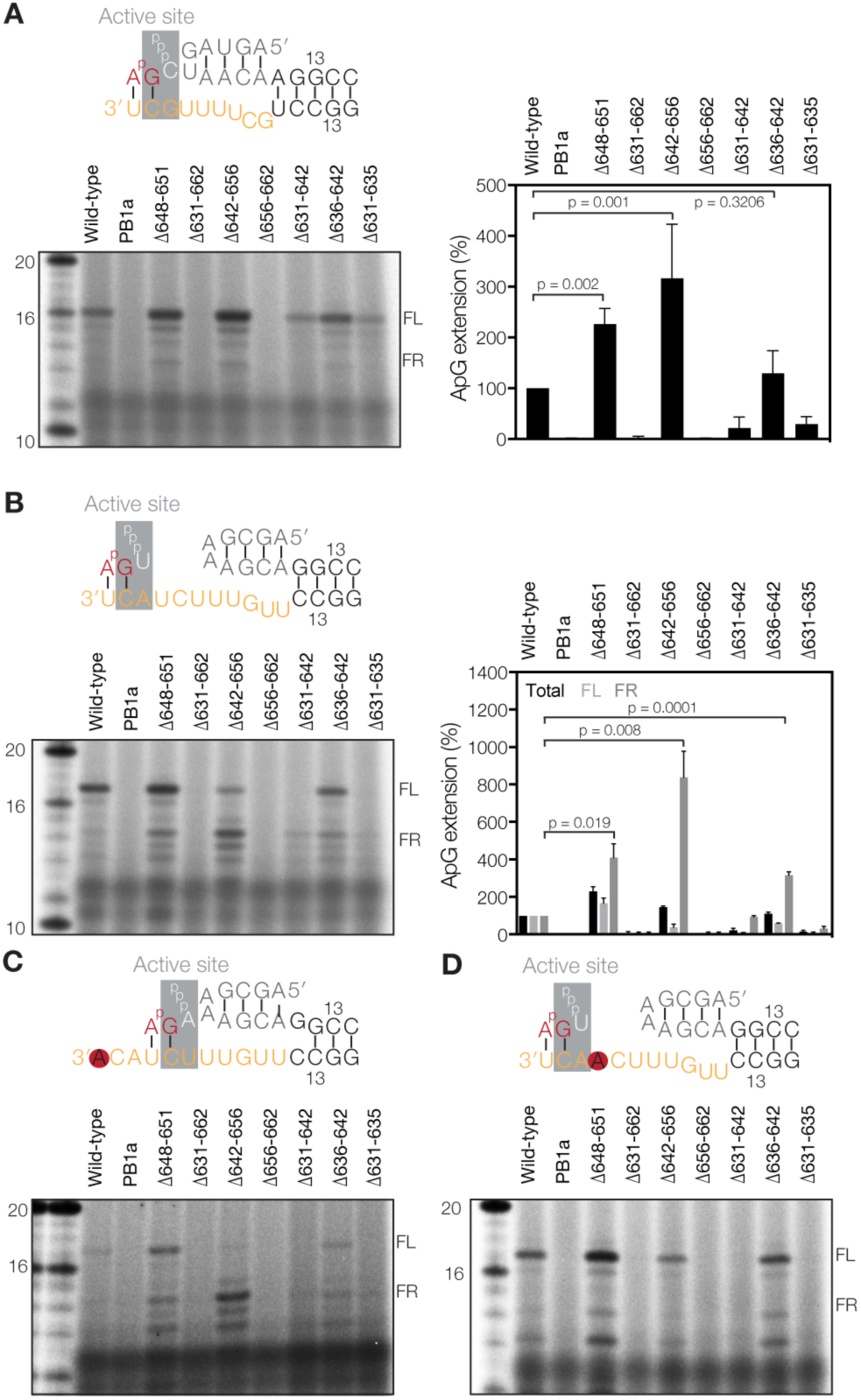
Deletions in the priming loop affect viral RNA synthesis in cell culture. (**A**) Position of the priming loop relative to the active site, the PB1 helix-turn-helix motif that contains V273 (V273 helix) and the promoter binding pocket of the influenza RdRp (PDB 5MSG). For clarity, only the right side of the RdRp is shown. (**B**) Superposed structures of the bat IAV priming loop (PDB 4WSB), influenza B virus priming loop (PDBs 5MSG, 4WSA, 4WRT and 5EPI) and the influenza C virus priming loop (PDB 5D98). The thickness of the backbone is scaled by β-factor. Deleted portions of the priming loop are shaded in greys and labelled a-e. (**C**) Alignment of the PB1 amino acid sequences that constitute the priming loop of the IAV A/WSN/33 (H1N1), influenza B virus B/Michigan/22687/09, and influenza C virus C/JJ/50 RdRps. Colours and secondary structure annotations as in Fig. 2B. Deletions in the priming loop are indicated with dotted lines. Labels are based on Fig. 4B. (**D**) Analysis of the steady state segment 6 viral RNA levels. The graph shows the mean RNA levels with error bars indicating standard deviation (*n* = 3). The PB1a signal was subtracted as background. (**E**) SDS-PAGE of recombinant influenza A virus RdRps purified from 293T cells followed by silver staining or Western-blot analysis of PB1 and PA in recombinant RdRps. Bottom two panels show expression of PB1 and PA RdRp subunits in 293T cells and a tubulin loading control. (**F**) Schematic of the smFRET promoter binding assay. Conformations of the influenza virus vRNA promoter before and after binding are shown as well as the distance between the Atto647N (red) and Cy3 (orange) dyes on the two promoter strands. (**G**) vRNA promoter binding by the IAV RdRp as analysed by smFRET and fitting with a single Gaussian. The mean apparent FRET (E*) of the RNA only signal is indicated as a dotted line in each graph for reference. (**H**) Cleavage of a radiolabelled capped 20-nucleotide long RNA. Alternative cleavage products are indicated with an asterisk.

### Purification of priming loop mutants

The RNP reconstitution assay described above confirmed that the PB1 priming loop is important for viral RNA synthesis, but due to the interdependence of these activities in cell culture we were not able to observe effects on transcription or replication. To study replication in more detail, the wild-type RdRp, the PB1a mutant, and our seven deletion mutants were expressed in 293T cells and purified by IgG chromatography. SDS-PAGE and Western blot analysis of the purified proteins showed that eight of the nine recombinant enzymes were able to form heterotrimers (Fig. 4E). The only exception was mutant Δ631-662, for which the PB1 subunit appeared slightly underrepresented in the trimer and the PB1 Western-blot signal (Fig. 4E).

To investigate whether the Δ631-662 priming loop deletion affected RdRp-promoter complex formation, we used a single-molecule Förster resonance energy transfer (sm-FRET)-based binding assay (13, 24) (Fig. 4F). The FRET distribution of the fluorescently labelled RNA in solution resulted in an apparent FRET population with a mean and standard deviation of *E*\*=0.76±0.09 (Fig. 4G), whereas addition of the wild-type RdRp resulted in a shift to a lower apparent FRET due to a change in the RNA structure upon binding (Fig. 4F). By contrast, incubation with a promoter-binding mutant (Tetra mutant) (13) resulted in no shift in the apparent FRET population (Fig. 4G), demonstrating the specificity of the assay. When we next incubated the fluorescently labelled promoter with mutant Δ632-662 we observed a shift in the FRET population similar to the wild-type RdRp (Fig. 4G), suggesting that deletion of the priming loop had not affected promoter binding.

Next, we assessed whether the conformational change that the cap binding, endonuclease and 627 domains undergo upon promoter binding (1, 7) was impaired by the priming loop truncations. Since this rearrangement is crucial for cap cleavage, we incubated the priming loop mutants with a 20-nt long radiolabelled capped RNA. We observed that the activities of Δ631-662, Δ656-662, Δ631-642 and Δ631-635 were greatly impaired (Fig. 4H, Table 2). By contrast, the activity of mutants Δ648-651, Δ642-656 and Δ636-642 was indistinguishable from wild-type (Fig. 4H). Together, these controls suggest that mutants Δ648-651, Δ642-656 and Δ636-642 were folded, binding the viral promoter correctly and active, while the other four mutants had an unknown impairment that frustrated either cap cleavage or the conformational rearrangement of the RdRp domains.

**Table 2:**
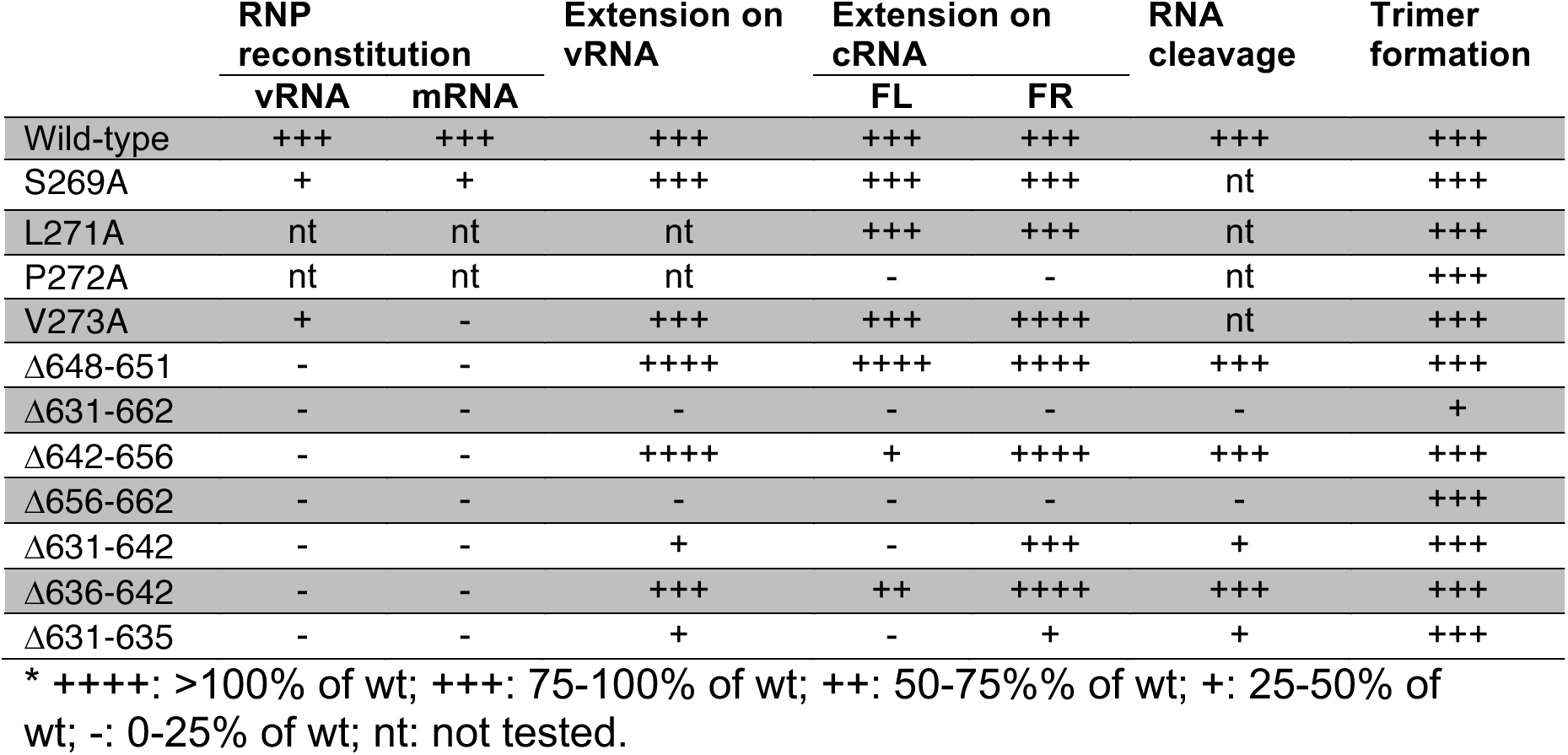
Overview of PB1 mutant characteristics relative to wild-type.

### The priming loop is important for prime-realignment

To investigate the effect of the priming loop deletions on the elongation of the RdRp, we first analysed the activity of the mutants in the presence of the vRNA promoter. Mutants Δ648-651 and Δ642-656 exhibited a higher activity compared to the wild-type enzyme, while the activity of the Δ636-642 mutant was indistinguishable from wild-type (Fig. 5A). In all three reactions, no differential change in the product pattern was observed (Fig. 5A). The four remaining priming loop mutants showed greatly impaired activities compared to wild-type (Fig. 5A).

**Figure 5.**
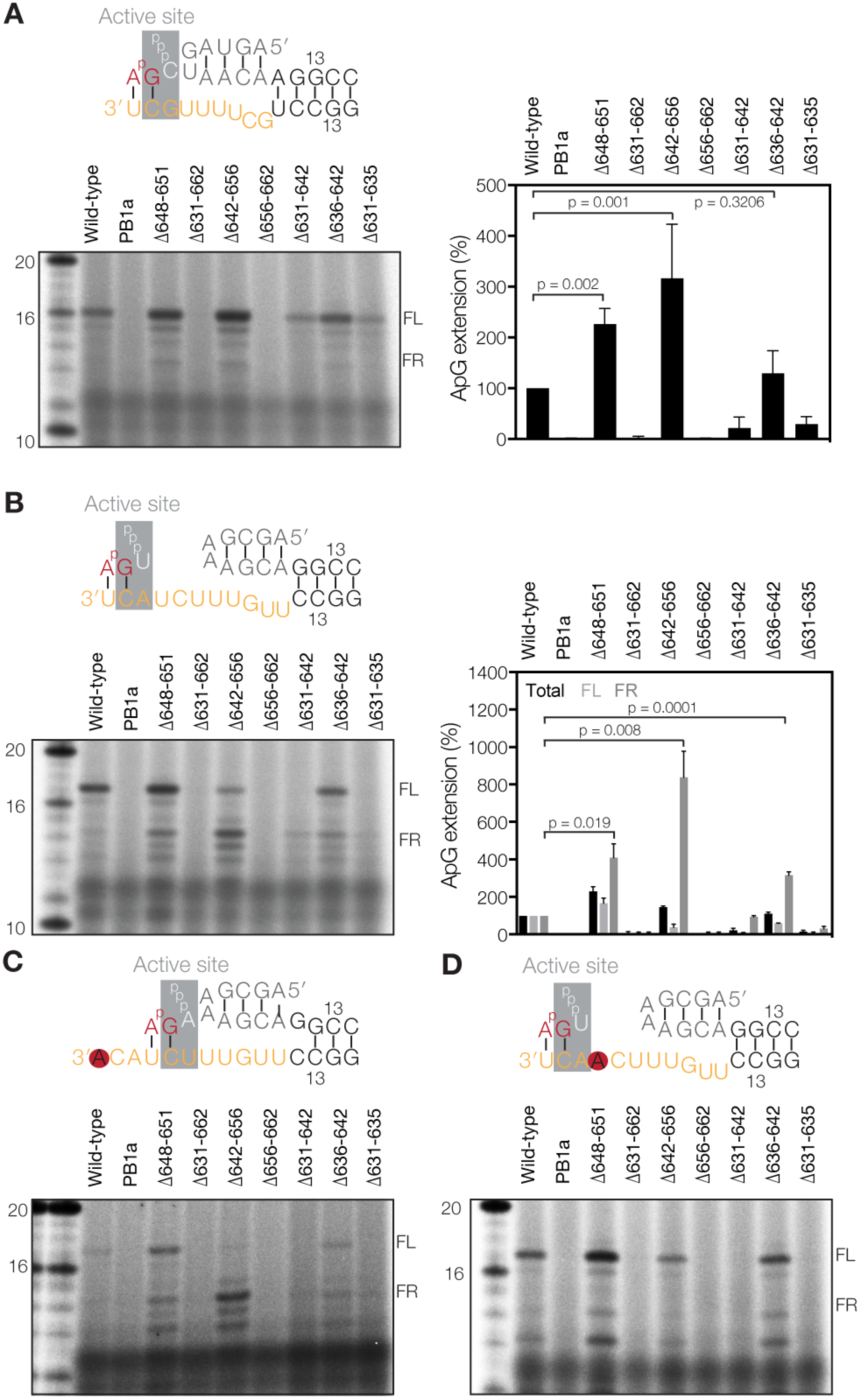
Deletions in the priming loop affect realignment *in vitro*. (**A**) ApG extension on the vRNA promoter and quantitation of the total vRNA extension activity relative to wild-type. (**B**) ApG extension on a cRNA promoter. Graph shows quantitation of the total ApG extension activity and the production of individual FR or FL bands relative to wild-type. (**C**) ApG extension on the mutant cRNA promoter 1U→A. (**D**) ApG extension on the mutant cRNA promoter 4U→A. Graphs show mean activity with error bars indicating standard deviation (*n* = 3). In Fig. 4A and Fig. 4B, p-values were determined using an unpaired t-test.

We next tested the effect of the mutations on prime-realignment by performing ApG extension assays on the cRNA promoter. We observed that mutants Δ648-651 and Δ642-656 exhibited a higher total activity compared to the wild-type enzyme, but that this signal contained more incorrectly realigned RNA as shown by the significantly stronger FR band (Fig. 5B). In the case of mutant Δ636-642, the total activity was composed of a strong increase in FR product synthesis and a reduction in FL product formation. The activity of the remaining four priming loop mutants was again greatly impaired (Fig. 5B), in line with the effect on cap cleavage (Fig. 4H) and ApG extension on the vRNA promoter (Fig. 5A).

To verify that the observed FR product was synthesised due to a failure in the prime-realign mechanism, we replaced the wild-type cRNA promoter with either the 1U→A promoter to reduce realignment (Fig. 5C) or the 4U→A promoter to prevent internal elongation (Fig. 5D). We found that mutants Δ648-651, Δ642-656, and Δ636-642 were still able to produce the incorrectly realigned RNA on the 1U→A promoter (Fig. 5C). By contrast, on the 4U→A promoter the Δ642-656 mutant showed a dramatic decrease in the FR band, whereas the Δ648-651 and Δ636-642 mutants still produced some FR signal (Fig. 5D). Interestingly, mutant Δ648-651 was also able to produce an FL product on the 1U→A promoter, while the other mutants and the wild-type polymerase were not, suggesting that this mutant had an increased tolerance for mismatches between the template and the dinucleotide primer (Fig. 5C). Overall, these results are consistent with a model in which the priming loop stimulates realignment during IAV vRNA synthesis and plays a role in suppressing internal elongation, thereby contributing to correct vRNA synthesis.

## Discussion

The two initiation mechanisms that drive IAV replication are substantially different. *De novo* initiation on a vRNA promoter occurs on the terminus of the template and critically depends on the PB1 priming loop (Fig. 1B and 6) (13). By contrast, *de novo* initiation on the cRNA promoter uses internal residues as template (13) (Fig. 1B and 6) and requires a realignment step to translocate the nascent vRNA to the terminus of the cRNA promoter before extension can take place. Here we show that a failure to efficiently perform this realignment step generates vRNAs lacking the first 3 nucleotides of the canonical vRNA 5ʹ terminus, which in turn impairs viral cRNA and mRNA synthesis in cell culture (Fig. 3, Table 2). In addition, we provide insight into the mechanism controlling this process by demonstrating that the priming loop plays a critical role in the efficiency of the realignment mechanism (Table 2). Based on observations from other RNA virus RdRps, in which the priming loop must undergo a conformational change to allow the product duplex to leave the active site (25, 26), we propose that the priming loop acts as ‘a spring’ that needs to undergo a conformational change to allow efficient elongation of the nascent replication product (Fig. 6). Thus, when the nascent vRNA is short and bound internally on the cRNA, the nascent vRNA cannot induce a conformational change in the priming loop. As a result, the nascent vRNA is destabilised, enabling realignment. However, when the duplex is extended to at least 4 nt, which can only occur when terminal elongation takes place, a conformational change in the priming loop can be induced, allowing the nascent RNA to exit the active site (Fig. 6).

**Figure 6.**
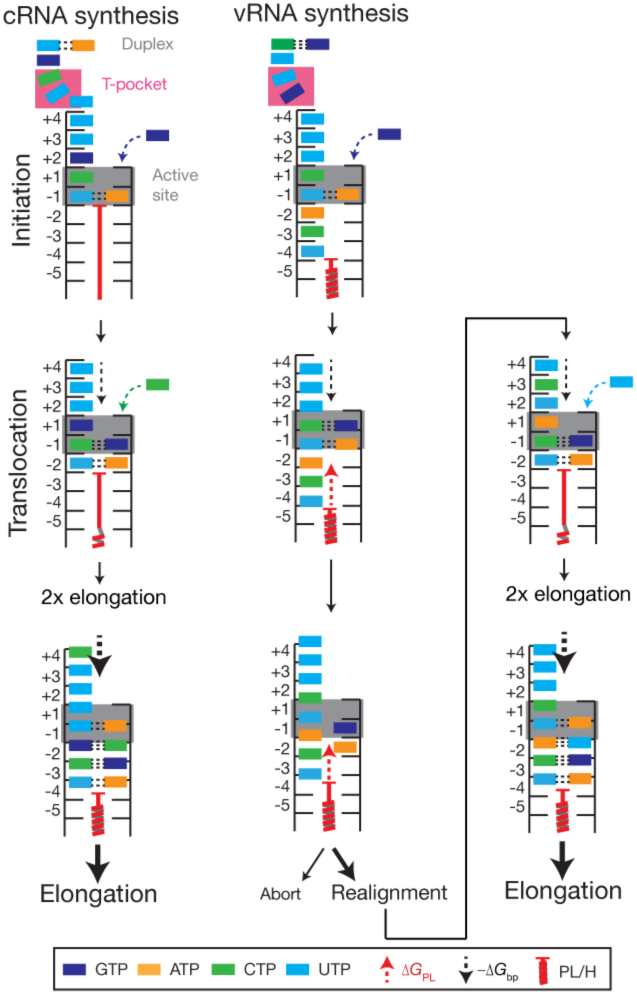
Model of influenza A virus replication initiation. cRNA synthesis is initiated on 3ʹ 1U and 2C of the vRNA when these occupy position −1 and +1 of the active site. Residues 7U and 8C are stacked in a T-orientation by residues of the PB2 subunit (the ‘T-pocket’), which likely fixes this end of the vRNA in the RdRp. Indeed, it is likely that the interaction between PB2 and 7U is sequence/structure specific as a 7U→A mutation was previously shown to abrogate *de novo* initiation (14). To stabilise the initiating ATP, an interaction with the tip of the priming loop is required. Translocation is not hindered by the priming loop after initiation, allowing two elongation steps until a tetramer is formed. Then a further NTP incorporation induces a conformational change in the priming loop, potentially similar to HCV elongation (25), in order to facilitate progressive elongation. The stability of the template-product duplex (-Δ*G*_bp_) is high enough to overcome a translocation block by the priming loop (Δ*G*_PL_). By contrast, initiation of a nascent vRNA is catalysed using cRNA residues 3ʹ 4U and 5C as template when these occupy position −1 and +1 of the active site. Although no structure is available of the RdRp bound to the complete cRNA promoter, it is plausible that G9 will occupy the T-pocket when the cRNA 3ʹ end enters the template channel, fixing one end of the cRNA in place. Indeed, previous mutational data have shown that like 7U in the vRNA, G9 in the cRNA is essential for *de novo* initiation (14). After dinucleotide formation, the priming loop blocks translocation of 3ʹ 1U to position −5, which induces melting of the template-dinucleotide duplex. The cRNA 3ʹ terminus can now move back 2 positions until 3ʹ 1U occupies position −2 to facilitate hybridisation with the dinucleotide and allow extension. vRNA extension now becomes effectively similar to cRNA synthesis. The influences of the priming loop/palm subdomain helix (PL/H) structures are drawn as a single red coiled structure for simplicity. The probability of realignment or elongation is indicated by the font size.

The above process relies on the correct positioning of the cRNA terminus in the active site. In a recent influenza B virus transcription initiation structure, the vRNA 3ʹ strand was shown to enter the template entry channel with 6 nt (17) (Fig. 4A), overshooting positions −1 and +1 of the active site by 1 nt. From this position, the template must move back 1 nt to allow terminal *de novo* initiation (Fig. 6). By contrast, to support internal *de novo* initiation, 8 nt must enter the template entry channel (Fig. 6). The IAV RdRp can readily achieve this for the cRNA template, because the cRNA 3ʹ promoter strand is one nt longer than the vRNA 3ʹ strand and the promoter duplex 1 nt shorter (Fig. 1B and 6). Moreover, our data show that reducing the stability of the cRNA promoter duplex does not increase initiation on a cRNA promoter (Fig. 1). Together, these different lines of research suggest that 4U and C5 of the cRNA 3ʹ promoter strand can be placed at positions −1 and +1 of the active site without duplex unwinding.

After internal *de novo* initiation, the dinucleotide-template duplex melts and the 3ʹ terminus of the cRNA templatemust move back to allow the pppApG dinucleotide to rebind at 1U and 2C in positions −1 and +1 of the active site (Fig. 6). We have previously reported that the 3ʹ end of the cRNA promoter can move freely in and out of the template channel using single-molecule experiments (12). We here add evidence that the priming loop and palm subdomain residue V273 may stimulate this process (Fig. 2 and 5) by acting as elongation block (Fig. 6). Importantly, the priming loop does not induce realignment after terminal initiation on a vRNA promoter (Fig. 5), which is consistent with the idea that the stability of the vRNA template-tetramer duplex is sufficient to prevent duplex melting and induce a conformational change in the priming loop (Fig. 6).

In the absence of a structure of the influenza virus RdRp elongation complex, we can only speculate about the mechanism through which V273 affects realignment. Interestingly, in both the apo structure of the influenza C virus RdRp (7) and the influenza B virus RdRp bound to the cRNA 5ʹ terminus (11), the PB2 cap binding domain closes the nascent strand exit channel via interactions with the PB1 helix-turn-helix, in which V273 resides, and the PB2 mid-domain (1, 7). If the conformation of the RdRp that has bound the cRNA 5ʹ terminus is indeed representative of the replicative form of the IAV RdRp, the PB2 cap binding domain must undergo a conformational change to allow the nascent strand to exit the polymerase. In support of this idea, a recent study showed that deletion of the cap binding domain affects vRNA synthesis, but not cRNA synthesis (27) and it is thus tempting to speculate that V273 may be involved in or positioned through the packing of the PB2 cap binding domain against the template exit channel and thereby indirectly affecting elongation and realignment. Future studies are needed to address this model in more detail.

In summary, we have here analysed how the IAV RdRp controls realignment during viral replication, a process that is a critical step in influenza virus RNA synthesis. Our findings offer new insights into influenza virus replication and they may have implications for the many other RNA viruses that also rely on a prime-realign mechanism to replicate and/or transcribe their genome (16, 28–30).

## Experimental procedures

### Cells and plasmids

Human embryonic kidney (HEK) 293T cells were mycoplasma tested and maintained in DMEM (Sigma) supplemented with 10% fetal calf serum (FCS). Plasmids pPolI-NA, pcDNA-NP, pcDNA-PB1, pcDNA-PA, pcDNA-PB2-TAP, and pcDNA-PB1a have been described previously (31–33). Also the promoter binding mutant (PB1 R238A, R672A and PA K572A and K583A; or “Tetra mutant”) (13), the priming loop mutant PB1 Δ648-651 (13), the PA endonuclease mutant D108A (34), and the PB1 palm subdomain mutants S269A and V273A have been reported before (19). To construct plasmids expressing additional mutant forms of the PB1 subunit, the plasmid pcDNA-PB1 was altered using side-directed mutagenesis with the primers (Life Technologies) listed in Table 1.

### Sequence alignment and structural modelling

Amino acid sequences of the PB1 subunits of IAV A/WSN/33 (H1N1), influenza B virus B/Michigan/22687/09, and influenza C virus C/JJ/50 were aligned using ClustalX (35) and visualised using ESPript (36). To visualise the influenza B virus RdRp crystal structure (PDB 5MSG) Pymol 1.3 was used. To model the poliovirus 3D^pol^ elongation complex (PDB 3OL7) into the influenza B virus RdRp crystal structure (PDB 5MSG), we aligned active site residues 324-332 of the poliovirus enzyme with residues 442-449 of the influenza virus PB1 protein in Pymol 1.3.

### Purification of recombinant influenza virus RNA polymerase

Wild-type and mutant recombinant RdRp preparations were purified using tap affinity purification (TAP) tags on the C-terminus of the PB2 subunit (33). The recombinant polymerases were expressed via transfection of 3 µg of PB1 or mutant PB1, PB2-TAP, and PA into HEK 293T cells using Lipofectamine^2000^ (Invitrogen). After 48 h the cells were lysed and the recombinant protein purified using IgG Sepharose (GE Healthcare) chromatography and cleavage by tobacco etch virus (TEV) protease (Invitrogen) in cleavage buffer (20 mM Hepes pH 7.5, 150 mM NaCl, 0.1% NP40, 10% glycerol, 1x PMSF and 1 mM DTT). The recombinant RdRp preparations were analysed by 8% SDS-PAGE and silver staining using a SilverXpress kit (Invitrogen). The concentration of the proteins was estimated in gel using a BSA standard. For Western-blot, polyclonal PB1 (GTX125923, Genetex), PB2 (GTX125926, Genetex) and PA (GTX118991, Genetex) antibodies were used. For activity assays, the RdRp preparations were stored in cleavage buffer at −80 °C. RdRp preparations for smFRET experiments were concentrated in 50 mM Hepes (pH 7.5), 10% glycerol, 500 mM NaCl, 0.05% *n*-Octyl β-D-thioglucopyranoside (OTG), and 0.5 mM tris(2-carboxyethyl)phosphine (TCEP) before storage at −80 °C.

### RNP reconstitution assay

RNP reconstitutions were performed by transfecting plasmids expressing PB1 or mutant PB1, PA, PB2, and NP together with pPolI-NA into HEK 293T cells using Lipofectamine^2000^ (Invitrogen). Typically, 0.25 *µ*g of each plasmid was used for a 6-well transfection. Total RNA was isolated using Tri Reagent (Sigma) 24 h after transfection and analysed using primer extension assays as described previously (13). Briefly, viral RNA species and a 5S rRNA loading control were reverse transcribed using ^32^P-labelled primers 1280NA, NA160 and 5S100 (Table 1), and SuperScript III (Invitrogen). cDNA synthesis was stopped with 10 μl loading buffer (90% formamide, 10% ddH_2_O, 10 μM EDTA, xylene cyanole, bromophenol blue) and analysed by 6% denaturing PAGE (6% 19:1 acrylamide/bis-acrylamide, 1x TBE buffer, 7 M urea). The viral RNA species and the 5S rRNA signal were visualised using a FLA-500 scanner (Fuji) and analysed using AIDA (RayTek).

### *In vitro* ApG extension assay

ApG extension assays were performed as described previously (13). Briefly, 4-μl reactions were performed that contained: 500 μM ApG (Jena Bioscience), 1 mM DTT, 500 μM UTP (unless indicated otherwise), 500 μM CTP, 500 μM ATP (unless indicated otherwise), 0.7 μM vRNA or cRNA promoter (Sigma), 5 mM MgCl_2_, 1 U μl^−1^ RNAsin (Promega), 0.05 μM [α-^32^P]GTP (3000 Ci mmole^−1^, Perking-Elmer), 5% glycerol, 0.05% NP-40, 75 mM NaCl, 10 mM HEPES pH 7.5, and ~2 ng RdRp µl^−1^. The reactions were incubated for 1 h at 30 °C and stopped with 4 μl loading buffer. The RNA products were analysed by 20% denaturing PAGE and visualised by phosphorimaging. P-values were determined using an unpaired non-parametric t-test.

### Capping of RNA primer and capped oligo cleavage assay

A synthetic 5’ triphosphate-containing 20 nt long RNA (ppAAUCUAUAAUAGCAUUAUCC, Chemgenes) was capped with a radiolabelled cap-1 structure using 0.25 µM [α-^32^P]GTP (3,000 Ci mmole^−1^, Perkin-Elmer), 2.5 U/µl 2’-O-methyltransferase (NEB) and a vaccinia virus capping kit (NEB) according the manufacturer’s instructions. The product was analysed by 20% denaturing PAGE, excised from the gel and desalted using NAP-10 columns (GE Healthcare) that had been equilibrated with RNase free water. To test the endonuclease activity of the IAV RdRp, we performed 3-μl reactions that contained: 1 mM DTT, 0.7 μM vRNA promotor (Sigma), 5 mM MgCl_2_, 1 U μl^−1^ RNAsin (Promega), 1500 cpm capped 20-nucleotide long RNA primer, 5% glycerol, 0.05% NP-40, 75 mM NaCl, 10 mM HEPES pH 7.5, and ~2 ng RdRp µl^−1^. The reactions were incubated for 1 hour at 30 °C, stopped with 4 μl loading buffer and analysed by 20% denaturing PAGE. The capped RNA cleavage products were visualised by phosphorimaging.

### *In vitro* dinucleotide synthesis assay

The pppApG synthesis activity was measured as described previously (13). Briefly, 3-μl reactions were set up that contained: 1 mM DTT, 350 μM adenosine, 5 mM MgCl_2_, 1U μl^−1^ RNAsin, 0.05 μM [α-^32^P] GTP (3000 Ci/mmole, Perking-Elmer), 0.7 μM vRNA or cRNA promotor (Sigma), 5% glycerol, 0.05% NP-40, 75 mM NaCl, 10 mM HEPES pH 7.5, and ~2 ng RdRp µl^−1^. The reactions were incubated for 18 h at 30°C, inactivated for 2 min at 95 °C and then treated with 1 U calf intestine alkaline phosphatase (Promega) at 37°C for 30 min. The reactions were stopped with 4 μl loading buffer and analysed by 20% denaturing PAGE.

### Single-molecule Förster resonance energy transfer

Promoter binding was measured as described previously (12, 13, 24). Briefly, a Cy3 donor dye was placed on 17U of the 3ʹ promoter strand and an Atto647N acceptor dye was placed on 6U of the 5ʹ strand. The RNA oligonucleotides were synthesized by IBA, and labelled, purified, and annealed as described previously (24). The excitation of the donor and acceptor fluorophores was measured using a custom-built confocal microscope with alternating-laser excitation (ALEX) (37, 38). In a typical experiment, ~100 nM RdRp was pre-incubated with 1 nM double-labelled promoter RNA in binding buffer (50 mM Tris-HCl (pH 8.0), 5% glycerol, 500 mM NaCl, 10 mM MgCl_2_, 100 µg/ml BSA) for 15 min at 28 °C. Samples were diluted 10-fold in binding buffer before the measurements were performed at excitation intensities of 250 µW at 532 nm and 60 µW at 635 nm. The *E\** values were plotted as one-dimensional distributions and fitted with a single Gaussian to obtain the mean *E\** and the standard deviation.

## Acknowledgements

The authors thank Dr Ervin Fodor, Dr. Nicole Robb, Dr. Achilles Kapanidis and Benjamin Nilsson for support and discussions. This work was funded by Wellcome Trust grants 098721/Z/12/Z and 206579/Z/17/Z (to AJWtV), grant 825.11.029 from the Netherlands Organization for Scientific Research (to AJWtV) and an Erasmus+ mobility grant (to JO).

## Author contributions

AJWtV designed experiments. JO and AJWtV performed experiments and analysed data. AJWtV wrote manuscript with input from JO.

## References

1. te Velthuis AJ, Fodor E. Influenza virus RNA polymerase: insights into the mechanisms of viral RNA synthesis. 2016. Nat Rev Microbiol 18:473–493

2. Fodor E. The RNA polymerase of influenza a virus: mechanisms of viral transcription and replication. 2013. Acta Virol 57:113–122

3. York A, Hengrung N, Vreede FT, Huiskonen JT, Fodor E. Isolation and characterization of the positive-sense replicative intermediate of a negative-strand RNA virus. 2013. Proc Natl Acad Sci U S A 110:E4238–E4245

4. Pflug A, Lukarska M, Resa-Infante P, Reich S, Cusack S. Structural insights into RNA synthesis by the influenza virus transcription-replication machine. 2017. Virus Research S0168-1702:30782–30781

5. Reich S, Guilligay D, Pflug A, Malet H, Berger I, Crépin T, Hart D, Lunardi T, Nanao M, Ruigrok RW, Cusack S. Structural insight into cap-snatching and RNA synthesis by influenza polymerase. 2014. Nature 516:361–366

6. Pflug A, Guilligay D, Reich S, Cusack S. Structure of influenza A polymerase bound to the viral RNA promoter. 2014. Nature 516:355–360

7. Hengrung N, El Omari K, Serna Martin I, Vreede FT, Cusack S, Rambo RP, Vonrhein C, Bricogne G, Stuart DI, Grimes JM, Fodor E. Crystal structure of the RNA-dependent RNA polymerase from influenza C virus. 2015. Nature 527:114–117

8. Parvin JD, Palese P, Honda A, Ishihama A, Krystal M. Promoter analysis of influenza virus RNA polymerase. 1989. Journal of virology 63:5142–5152

9. Pritlove DC, Fodor E, Seong BL, Brownlee GG. In vitro transcription and polymerase binding studies of the termini of influenza A virus cRNA: evidence for a cRNA panhandle. 1995. J Gen Virol 76:2205–2213

10. Fodor E, Pritlove DC, Brownlee GG. The influenza virus panhandle is involved in the initiation of transcription. 1994. J Virol 68:4092–4096

11. Thierry E, Guilligay D, Kosinski J, Bock T, Gaudon S, Round A, Pflug A, Hengrung N, El Omari K, Baudin F, Hart DJ, Beck M, Cusack S. Influenza Polymerase Can Adopt an Alternative Configuration Involving a Radical Repacking of PB2 Domains. 2016. Molecular Cell 61:1–13

12. Robb NC, te Velthuis AJ, Wieneke R, Tampé R, Cordes T, Fodor E, Kapanidis AN. Single-molecule FRET reveals the pre-initiation and initiation conformations of influenza virus promoter RNA. 2016. Nucleic Acids Res 44:10304–10315

13. te Velthuis A, Robb NC, Kapanidis AF, Fodor F. The role of the priming loop in influenza A virus RNA synthesis. 2016. Nature Microbiology 1:16029

14. Deng T, Vreede FT, Brownlee GG. Different de novo initiation strategies are used by influenza virus RNA polymerase on its cRNA and viral RNA promoters during viral RNA replication. 2006. J Virol 80:2337–2348

15. Martin A, Hoefs N, Tadewaldt J, Staeheli P, Schneider U. Genomic RNAs of Borna disease virus are elongated on internal template motifs after realignment of the 3’ termini. 2011. Proc Natl Acad Sci U S A 108:7206–7211

16. Garcin D, Lezzi M, Dobbs M, Elliott RM, Schmaljohn C, Kang CY, Kolakofsky D. The 5’ ends of Hantaan virus (Bunyaviridae) RNAs suggest a prime-and-realign mechanism for the initiation of RNA synthesis. 1995. J Virol 69:5754–5762

17. Reich S, Guilligay D, Cusack S. An in vitro fluorescence based study of initiation of RNA synthesis by influenza B polymerase. 2017. Nucleic acids research 45:3353–3368

18. Vreede FT, Brownlee GG. Influenza virion-derived viral ribonucleoproteins synthesize both mRNA and cRNA in vitro. 2007. J Virol 81:2196–2204

19. Jung TE, Brownlee GG. A new promoter-binding site in the PB1 subunit of the influenza A virus polymerase. 2006. J Gen Virol 87:679–688

20. Du Y, Wu NC, Jiang L, Zhang T, Gong D, Shu S, Wu TT, Sun R. Annotating Protein Functional Residues by Coupling High-Throughput Fitness Profile and Homologous-Structure Analysis. 2016. MBio 7:e01801–16.

21. Laurila MR, Makeyev EV, Bamford DH. Bacteriophage phi 6 RNA-dependent RNA polymerase: molecular details of initiating nucleic acid synthesis without primer. 2002. J Biol Chem 277:17117–17124

22. Hong Z, Cameron CE, Walker MP, Castro C, Yao N, Lau JY, Zhong W. A novel mechanism to ensure terminal initiation by hepatitis C virus NS5B polymerase. 2001. Virology 285:6–11

23. Mosley RT, Edwards TE, Murakami E, Lam AM, Grice RL, Du J, Sofia MJ, Furman PA, Otto MJ. Structure of hepatitis C virus polymerase in complex with primer-template RNA. 2012. J Virol 86:6503–6511

24. Tomescu AI, Robb NC, Hengrung N, Fodor E, Kapanidis AN. Singlemolecule FRET reveals a corkscrew RNA structure for the polymerase-bound influenza virus promoter. 2014. Proc Natl Acad Sci USA 111:E3335–E3342

25. Appleby TC, Perry JK, Murakami E, Barauskas O, Feng J, Cho A, Fox D, Wetmore DR, McGrath ME, Ray AS, Sofia MJ, Swaminathan S, Edwards TE. Structural basis for RNA replication by the hepatitis C virus polymerase. 2015. Science 347:771–775

26. Tao Y, Farsetta DL, Nibert ML, Harrison SC. RNA synthesis in a cage— structural studies of reovirus polymerase lambda3. 2002. Cell 111:733–745

27. Nilsson BE, Te Velthuis AJ, Fodor E. Role of the PB2 627 Domain in Influenza A Virus Polymerase Function. 2017. J Virol 91: e02467–16

28. van Knippenberg I, Lamine M, Goldbach R, Kormelink R. Tomato spotted wilt virus transcriptase in vitro displays a preference for cap donors with multiple base complementarity to the viral template. 2005. Virology 335:122-130

29. Bouloy M, Pardigon N, Vialat P, Gerbaud S, Girard M. Characterization of the 5’ and 3’ ends of viral messenger RNAs isolated from BHK21 cells infected with Germiston virus (Bunyavirus). 1990. Virology 175:50–58

30. Duijsings D, Kormelink R, Goldbach R. In vivo analysis of the TSWV cap-snatching mechanism: single base complementarity and primer length requirements. 2001. EMBO J 20:2545–2552

31. Vreede FT, Jung TE, Brownlee GG. Model suggesting that replication of influenza virus is regulated by stabilization of replicative intermediates. 2004. J Virol 78:9568–9572

32. Fodor E, Crow M, Mingay LJ, Deng T, Sharps J, Fechter P, Brownlee GG. A single amino acid mutation in the PA subunit of the influenza virus RNA polymerase inhibits endonucleolytic cleavage of capped RNAs. 2002. J Virol 76:8989–9001

33. Deng T, Sharps J, Fodor E, Brownlee GG. In vitro assembly of PB2 with a PB1-PA dimer supports a new model of assembly of influenza A virus polymerase subunits into a functional trimeric complex. 2005. J Virol 79:8669-8674

34. Hara K, Schmidt FI, Crow M, Brownlee GG. Amino acid residues in the N-terminal region of the PA subunit of influenza A virus RNA polymerase play a critical role in protein stability, endonuclease activity, cap binding, and virion RNA promoter binding. 2006. J Virol 80:7789–7798

35. Larkin MA, Blackshields G, Brown NP, Chenna R, McGettigan PA, McWilliam H, Valentin F, Wallace IM, Wilm A, Lopez R, Thompson JD, Gibson TJ, Higgins DG. Clustal W and Clustal X version 2.0. 2007. Bioinformatics 23:2947–2948

36. Robert X, Gouet P. Deciphering key features in protein structures with the new ENDscript server. 2014. Nucleic Acids Res 42:W320–W324

37. Kapanidis AN, Lee NK, Laurence TA, Doose S, Margeat E, Weiss S. Fluorescence-aided molecule sorting: analysis of structure and interactions by alternating-laser excitation of single molecules. 2004. Proc Natl Acad Sci U S A 101:8936–8941

38. Santoso Y, Hwang LC, Le Reste L, Kapanidis AN. Red light, green light: probing single molecules using alternating-laser excitation. 2008. Biochem Soc Trans 36:738–744

39. te Velthuis AJW, Oymans J. Initiation, elongation and realignment during influenza virus mRNA synthesis. 2017. BioRxiv doi: https://doi.org/10.1101/19721000

